# A pan-gene catalogue of Asian cultivated rice

**DOI:** 10.1101/2025.02.17.638606

**Authors:** Bruno Contreras-Moreira, Eshan Sharma, Shradha Saraf, Guy Naamati, Parul Gupta, Justin Elser, Dmytro Chebotarov, Kapeel Chougule, Zhenyuan Lu, Sharon Wei, Andrew Olson, Ian Tsang, Disha Lodha, Yong Zhou, Zhichao Yu, Wen Zhao, Jianwei Zhang, Sandeep Amberkar, Kawinnat Sue-Ob, Maria Martin, Kenneth L. McNally, Doreen Ware, Zhi Sun, Eric W Deutsch, Dario Copetti, Rod A. Wing, Pankaj Jaiswal, Sarah Dyer, Andrew R Jones

## Abstract

The rice genome underpins fundamental research and breeding, but the Nipponbare (japonica) reference does not fully encompass the genetic diversity of Asian rice. To address this gap, the Rice Population Reference Panel (RPRP) was developed, comprising high-quality assemblies of 16 rice cultivars to represent *japonica*, *indica*, *aus*, and *aromatic* varietal groups.

The RPRP has been consistently annotated, supported by extensive experimental data and here we report the computational assignment, characterization and dissemination of stably identified pan-genes. We identified 25,178 core pan-genes shared across all cultivars, alongside cultivar-specific and family-enriched genes. Core genes exhibit higher gene expression and proteomic evidence, higher confidence protein domains and AlphaFold structures, while cultivar-specific genes were enriched for domains under selective breeding pressure, such as for disease resistance. This resource, integrated into public databases, enables researchers to explore genetic and functional diversity via a population-aware “reference guide” across rice genomes, advancing both basic and applied research.

## Introduction

Rice is one of the most important crops for human nutrition and will be central to efforts to feed 9.8 billion people by 2050. Breeding efforts are targeted with increasing yields and nutrition, while also making rice more resistant to biotic and abiotic stresses, which will be exacerbated by a changing climate. Asian cultivated rice *Orzya sativa* L. has previously been classified in different varietal groups, based on historic domestication events – mostly notably *indica* (*Xian*) and *japonica* (*Geng*), with other recognised groups including those described as *aus*, *aromatic* (Basmati) and *admixed*^1^.

An important resource underpinning both basic and applied rice research is the rice reference genome sequence, originally released via two draft assemblies circa 2002 (for *indica* and *japonica*)^2,3^, and then with one finished genome three years later by the International Rice Genome Sequencing Project (IRGSP), with an annotation set (gene models, gene and protein sequences) released in 2005^4^. The IRGSP canonical reference (IRGSP RefSeq) genome is derived from the *japonica* variety - Nipponbare, despite *indica* rice accounting for a much larger share of the international market. In the last 20 years, there have been multiple efforts to annotate the IRGSP reference, using different software packages and supporting data. From these efforts, two primary annotation sets have persisted – those developed by the Rice Annotation Project Database (RAP-DB)^5^ (example gene identifier: Os01g0918300), and the Rice Genome Annotation Project at Michigan State University^6^ (MSU) (example: LOC_Os01g68950). RAP-DB is still actively updating its database, while the MSU annotations have been frozen since 2014, with the latter being the more widely used/reference annotations (for historical reasons).

In order to study genetic variation, significant efforts have been made to identify variants on a large scale from high-throughput short DNA read sequencing, for example *via* the rice 3000 genomes project^1,7^. These projects can identify single nucleotide polymorphisms (SNPs) or short insertions and deletions, relative to the reference genome to which they are mapped. However, they cannot be used to find larger structural rearrangements of chromosomes, call variants relative to any regions absent from the reference genome used, or define accurate gene models for regions of chromosome absent from the reference genome.

Fully assembled genomes allow researchers to study chromosomal rearrangements and potentially identify gene gain or loss in different rice varieties. A collection of “platinum-standard” genomes assemblies, called the Rice Population Reference Panel (RPRP, and also called the “MAGIC-16”)^8,9^, has been created, including 15 new genomes (plus the IRGSP RefSeq) aimed specifically at covering a significant portion of the population genetic diversity of *Oryza sativa*. These genomes have now been annotated using a consistent pipeline (as described in Zhou *et al.*^8^), including support from long read transcriptome data in every variety, which is able to give strong experimental evidence for the correct splicing prediction of gene models. As new genomes are sequenced, assembled and annotated, they have the potential to act as a powerful resource for rice researchers, for example to find genes/variants present in only certain varieties, or where genes are differentially alternatively spliced across varieties. However, there are currently no stable/consistent nomenclature gene identifiers across the rice pan-genome, and it is not straightforward to determine the relationships between orthologous genes.

In this work, we built a pan-gene set for *Oryza sativa* from the annotated RPRP genomes, using an algorithm we developed based on whole genome alignment^10^. We assigned stable identifiers for a *pan-gene* set, here defined as a collection of gene models across different varieties in the same genomic location, following whole genome alignment i.e. syntenic orthologs. The members within each pan-gene could also be described as *Oryza sativa* alleles within different genomes. However, even with support from experimental data, accurate gene model definition remains highly challenging – i.e. genes adjacent on a chromosome may be falsely merged, a gene may be falsely split into two, splicing of exons may be incorrectly defined, and predicting the correct start codon is frequently wrong^11^. As such, some pan-genes will contain a mixture of some correct and some incorrect gene models, generated from the same (aligned) chromosomal region in different varieties. By bringing models together into assigned pan-genes, it allows work to begin to determine which gene models are correct and to refine annotations over time. Having defined pan-genes, we next explore their characteristics with respect to their occupancy across genomes, and the extent to which their gene expression and protein abundance is supported by experimental data from multiple sources. All gene annotation sets and pan-gene identifiers have been deposited into end-user focused databases – Ensembl Plants and Gramene (and protein sequences in UniProtKB), for straightforward user access.

## Results

### RPRP gene sets

We first collated gene models for the RPRP genomes, as a source for the creation of a pan-Oryza-gene set. All 16 genomes, including the IRGSP RefSeq were annotated using a consistent pipeline and data source (see Methods). In addition, there are also RAP-DB and MSU gene sets for the IRGSP RefSeq, making 18 annotation sets in total from which we built the pan-gene set. The count of gene models for each genome used as input to the pan-gene set are provided in Table 1. Around 36,000 genes were found in all varieties, with the exception of Nipponbare, which has a greatly inflated gene count, due to the merging of three input sets (See “nipponbare_merged_GFF” at https://zenodo.org/records/14772953).

**Table 1.** Summary information on the RPRP genomes and their annotations.

| Stock_Ac<br>cession | RPRP<br>gene<br>model<br>prefix | Genetic<br>Stock<br>Varname | Country_<br>Origin | K15_sub<br>pops | K15_com<br>mon | K5_grou<br>ps | Gene<br>count |
| --- | --- | --- | --- | --- | --- | --- | --- |
| IRGC<br>117425 | OsARC_ | ARC<br>10497::IR<br>GC<br>12485-1 | India | cB | cBasmati | Aro | 36442 |
|  | OsAzu_ | Azucena | Philippin<br>es | GJ-trop1 | tropA | Tropjap | 36626 |
| IRGC<br>132278 | OsCMeo<br>_ | CHAO<br>MEO::IR<br>GC<br>80273-1 | Lao PDR | GJ-subtrp | subtrop | Tropjap | 36999 |
| IRGC<br>132424 | OsGoSa_ | GOBOL<br>SAIL<br>(BALAM):<br>:IRGC<br>26624-2 | Banglade<br>sh | XI-2A | ind2A | indica | 36237 |
|  | OsIR64_ | IR 64 | Philippin<br>es | XI-1B1 | ind1B1 | indica | 36102 |
| IRGC<br>128077 | OsKeNa_ | KETAN<br>NANGKA:<br>:IRGC<br>19961-2 | Indonesi<br>a | GJ-trop2 | tropB | Tropjap | 36612 |
| IRGC<br>127518 | OsKYG_ | KHAO YAI<br>GUANG::<br>IRGC<br>65972-1 | Thailand | XI-3B1 | ind3B1 | indica | 36476 |
| IRGC<br>125619 | OsLaMu_ | LARHA<br>MUGAD::<br>IRGC<br>52339-1 | India | XI-2B | ind2B | indica | 36316 |
| IRGC<br>127564 | OsLima_ | LIMA::IR<br>GC<br>81487-1 | Indonesi<br>a | XI-3A | ind3A | indica | 37044 |
| IRGC<br>125827 | OsLiXu_ | LIU<br>XU::IRGC<br>109232-1 | China | XI-3B2 | ind3B2 | indica | 36378 |
|  | OsMH63<br>_ | Minghui<br>63 | China | XI-adm | ind-admx | indica | 36147 |
| IRGC<br>117534 | OsN22_ | N<br>22::IRGC<br>19379-1 | India | cA1 | cAus1<br>(Assam) | Aus | 36319 |
| IRGC<br>127652 | OsNaBo_ | NATEL<br>BORO::IR<br>GC<br>34749-1 | Banglade<br>sh | CA2 | cAus2<br>(Bengal) | Aus | 36196 |
|  | Os (RAP-DB);<br>Loc_Os<br>(MSU);<br>Gramene<br>annotati<br>ons<br>OsNip_ | Nipponb<br>are<br>(IRGSP1.<br>0) | Japan | GJ-temp | temp | Tempjap | 62225* |
| IRGC<br>127742 | OsPr106<br>_ | PR<br>106::IRG<br>C 53418-1 | India | XI-1B2 | ind1B2 | indica | 36436 |
|  | OsZS97_ | Zhen<br>Shan 97 | China | XI-1A | ind1A | indica | 35686 |

\*Breakdown by source, prior to merging: RAP-DB= 35,694, MSU=55,801 (including possible transposable elements), OsNip=37,007.

### Pan-gene sets, clusters and identifiers

Our first objective was to analyze gene annotation sets from the RPRP genomes and assign pan-genes following whole genome alignment (WGA), indicative of synteny. The GET_PANGENES pipeline was used to create pan-genes, determine their occupancy statistics and assign long-term stable identifiers (Figure 1). Supplementary table S1 contains the matrix of pan-genes, with their stable identifier (column 1), then 16 further columns - one per input genome, containing transcript identifiers (if any) from each genome that have been mapped to that pan-gene. There are a total of 77,530 pan-genes, including 35,029 singletons i.e. pan-genes containing transcripts from only one genome (17,383 singletons when MSU gene models are removed, which heavily inflate the count), leaving 42,501 pan-gene clusters containing two or more members. Figure 1A displays the distribution of pan-genes by occupancy class, demonstrating that most genes are core or cloud, and that shell genes are relatively rare in the pan-gene set.

**Figure 1.**
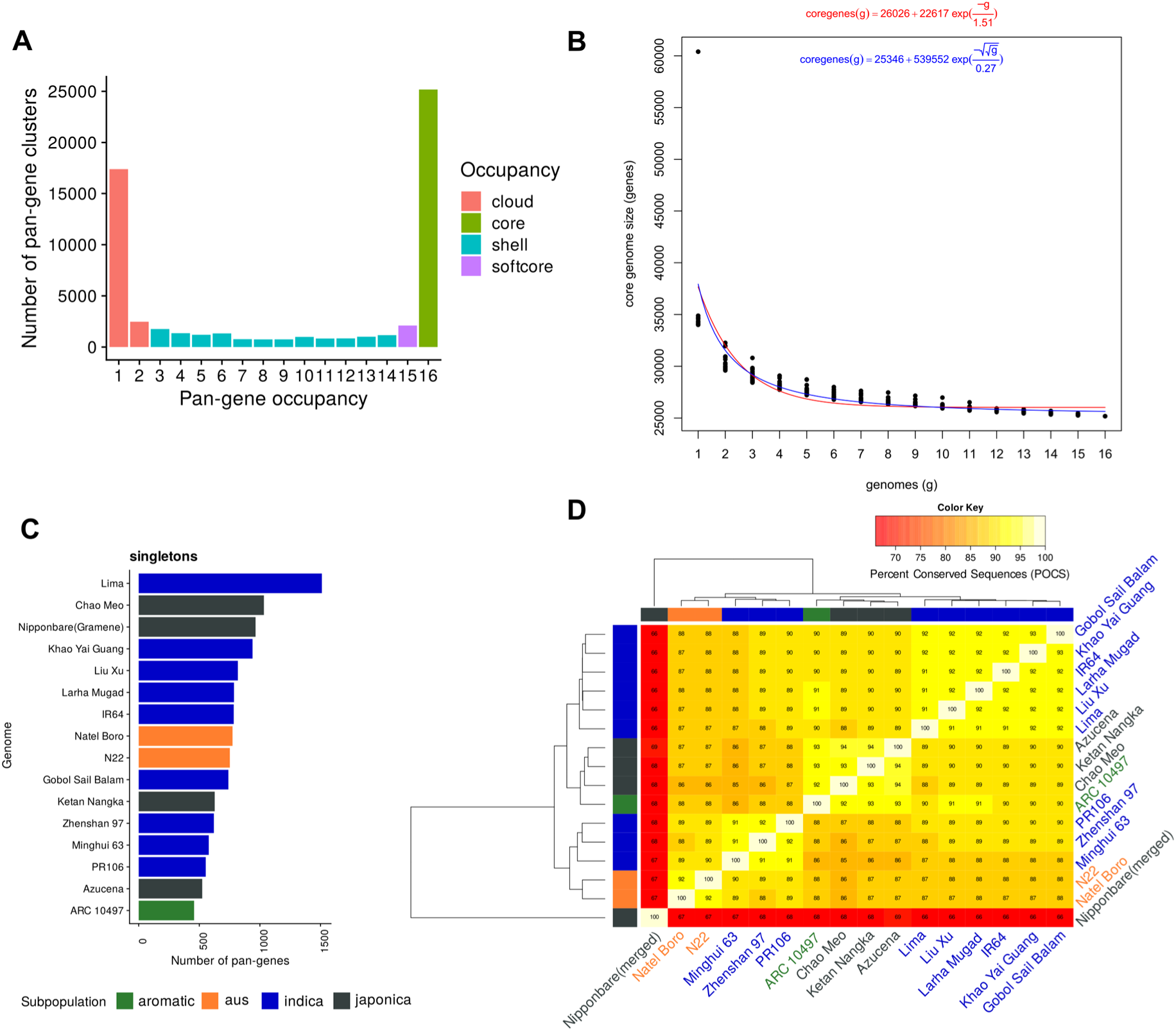
Pan-gene sets of rice MAGIC15 and Nipponbare cultivars. A) Count of input genome members (x-axis) “occupancy” 1-16 vs counts of pan-genes (pan-genes containing only one MSU gene model have been removed). B) Core-genome growth simulations after adding MAGIC15 cultivars in random order, using Tettelin^13^ (red) and Willenbrock^14^ (blue) functions, fitted after 20 permutation experiments. C) Counts of singletons i.e. genes per input genome present in clusters with no other gene models (for IRGSP, singleton count includes those with an OsNip (Gramene-generated) gene, for unbiased comparison); D) Matrix of % shared clusters and heat map of RPRP pan-genes. The dendrograms were computed by complete linkage clustering and Euclidean distances computed among columns.

We also assessed the quality and robustness of pan-gene formation through multiple metrics, including descriptive statistics on gene length, exon count, sequence distance between members, and “MSA completeness” (Extended data figure 1, Supplementary table S2), demonstrating that the majority of pan-genes contain highly similar gene / protein sequences from different genomes. The metrics also allow for identification of pan-genes containing one or more inconsistent members, indicative of different overlapping gene predictions in some genomes (e.g. coding sequences predicted on different frames), which are likely artifacts of incorrect gene model prediction.

Figure 1B demonstrates that the core set of genes converges at around 25,000 genes (24,931 and 23,651 via Tettelin and Willenbrock methods, respectively), indicating this is the minimally core set contained within an *Oryza sativa* genome. Figure 1C displays the counts per variety of singletons from each genome, ranging from 457 to 1511 (mean = 766, median 752), indicating consistency across different cultivars for the presence of these unique genes. For Nipponbare, the results are filtered to include singletons containing an OsNip-source model, created using the same approach as the other 15 genomes to avoid artefactually high counts due to merging different input gene sets, giving a set of 964 genes in line with other genomes. Figure 1D shows the pan-gene cluster overlap across cultivars. In general, there is reasonable clustering by rice family (*japonica* vs *aus* vs *indica*). The merged Nipponbare annotation has lower overlap with other clusters, due to it having a much higher count of input genes.

### Genomic position of pan-genes

In Extended data figure 2, we display the chromosomal location of genes within the pan-gene set, using a “pseudo-position” attribute (see Methods). There are sporadic cases of genes that are not collinear with respect to the reference genome, with a few notable systematic cases. These include a set of genes that are inverted on chromosome 1 for OsLima at position 600K-800K bp with respect to all other genomes. Similarly, there are two inversions on *aus* genomes (N22 and Natal Boro) around the center of chromosome 1. Another notable case is on chromosome 6, where Nipponbare has a large inversion compared to the other 15 genomes. Data containing the positions of genes within pan-gene clusters, and pseudo-positions of clusters can be found in Supplementary table S3. The GET_PANGENES approach uses whole genome alignment and does not enforce that genes must be located on the same chromosome. This enables the algorithm to identify cases where breeding (or natural selection) has caused regions of the chromosome to translocate, or potentially where there have been assembly errors. There are 1502 pan-genes that have occupancy > 2 and contain genes mapped to different chromosomes (Supplementary table S3), which researchers should consider when using genomics resources e.g. SNP mapping or allele mining.

### Pan-gene distribution across rice families

Supplementary table S4 contains the proportions of genomes by sub-family (*japonica*, *aromatic*, *aus*, *indica*) that have a member within each pan-gene. For example, by filtering this data, one can identify genes that segregate by family e.g. 359 pan-genes found in all Japonica genomes but only 0 or 1 other genome, 229 pan-genes found in 8 or 9 Indica genomes but present in 0 or 1 other genomes, 336 pan-genes found in Aus varieties and 0 or 1 other genomes. For multiple research and breeding purposes such genes are of high interest, particularly if there are QTLs mapped to these genomic regions. One of the current major limitations of genome wide association or other population SNP mapping studies is that the ISRFG RefSeq reference genome (IRGSP) is almost always used for read mapping, and locating genes nearby SNPs. Unfortunately, as we show here, there are 17,130 pan-genes that do not have an IRGSP gene model (5,634 with occupancy >=2), indicative of a large pool of genetic resources that is currently missed in most such studies.

### Properties of pan-genes by occupancy

We next explored the properties of pan-genes related to occupancy (Figure 2). We chose to focus on quality metrics derived from the RAP-DB reference gene annotations (for which most data already exist from different resources), and taking the occupancy statistic from the parent pan-gene. First, we assessed the quality of a given gene model, using the TRaCE^12^ algorithm, and assigned a minimum annotation edit distance (AED) score, from three transcriptomic data sets, to assess the quality of the annotation. AED is calculated on a scale of 0-1 based on the sequence distance between the gene model and the best transcript, assembled from publicly available transcriptomics data.

**Figure 2.**
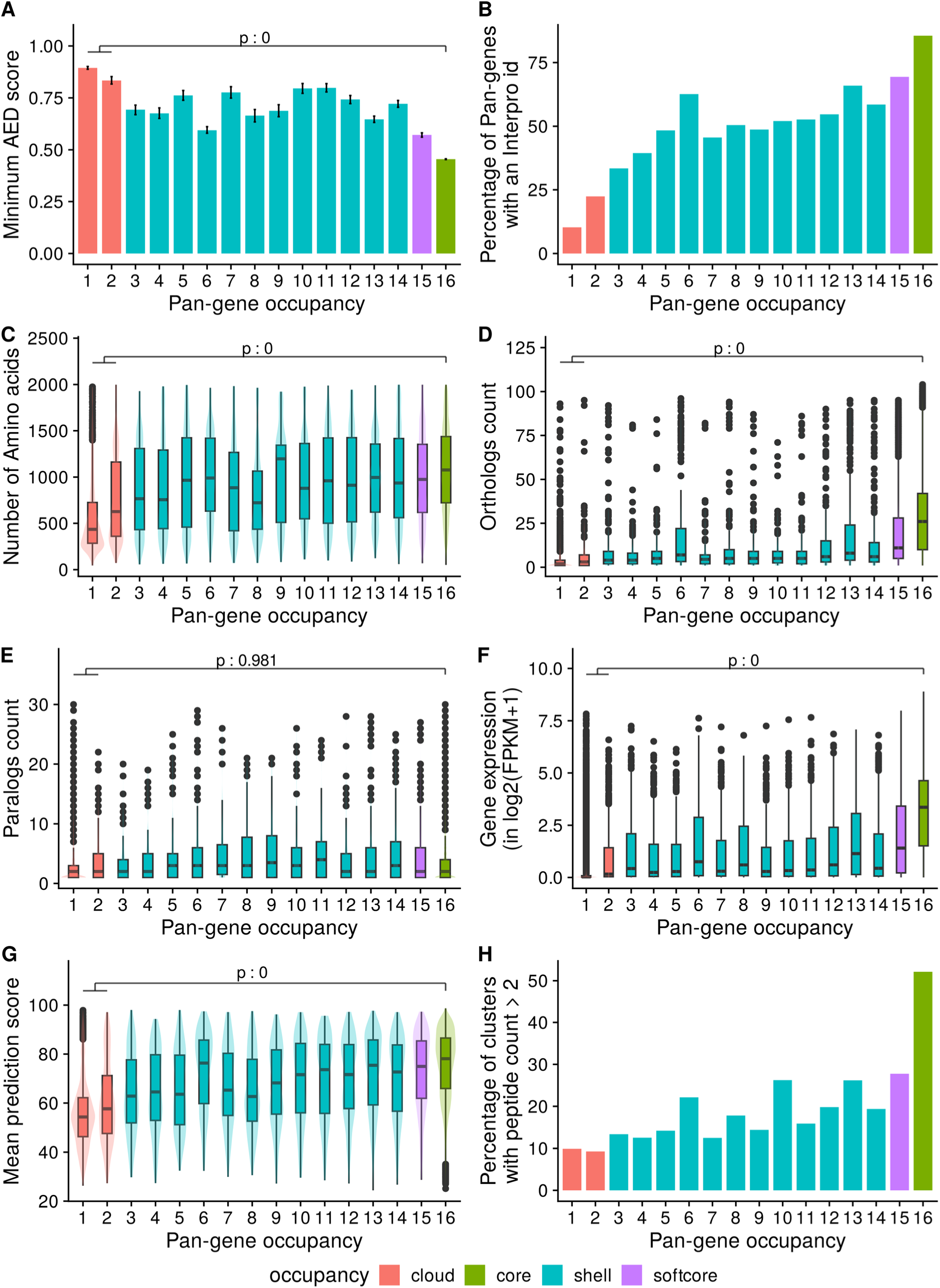
Exploration of pan-genes and their properties by their occupancy. The x-axis shows the pan-gene occupancy for each cluster, the y-axis shows data derived from RAP-DB gene models within those clusters (clusters containing no RAP-DB model are excluded): A) mean of minimum AED score from three transcriptomic data sets; B) proportion of genes that contain a significant match to an InterPro domain; C) Mean count of amino acids; D) boxplot showing the counts of orthologs derived from Ensembl Compara (see Methods); E) boxplot showing counts of paralogs; F) boxplot of mean quantitative gene expression data log2(FPKM+1), sourced from 11,726 rice RNA-Seq samples; G) Boxplot of mean prediction score (pLDDT) from corresponding AlphaFold2 model; H) Percentage of clusters with genes having peptide evidence (n >2 per gene).

Figure 2A shows the mean (minimum) AED score for all RAP-DB genes, against the occupancy of their parent pan-gene. We can observe that core (occupancy = 16) or softcore (15 or 16) clusters have significantly lower AED within core (mean=0.5) versus cloud pan-genes (mean=0.9) indicating that core genes are called with higher confidence using transcriptomic data sources. Of the core pan-gene, 83% have an assigned InterPro domain (Figure 2B), compared to 11% in singleton pan-genes (occupancy = 1). More detailed analysis of protein domains across all pan-genes is provided below. Core pan-genes are also significantly longer and are more likely conserved across the plant kingdom as indicated by their protein length and ortholog count (Figure 2C, D).

There is a more complex trend for paralog count (Figure 2E), where there is no significant difference between paralog counts for cloud or shell, versus core sets. There appears to be a trend where shell genes have higher paralog counts. We hypothesize that there are two “competing” distributions here. We would expect that pan-genes of lower occupancy are more likely to have paralogs, as this set contains rapidly duplicating gene families (further discussed below), but are also enriched for faulty gene models, where paralogs cannot be identified.

Pan-genes with softcore/core occupancy have significantly higher gene expression values compared to cloud genes, with singleton pan-genes (occupancy=1) frequently having no detectable signals – these could be pseudo-genes or genes only transcribed under particular conditions not assessed in the source data (Figure 2F). Prediction of a 3D protein structure for core/softcore and shell pan-genes can be done by AlphaFold with high confidence (Median <u>+</u> <u>IQR</u> score 78.1 ± 20.6 for core, 75.0 ± 23.4 for softcore and 70.3 ± 27.2 for shell pan-genes), whereas a protein structure for cloud pan-genes has a median score of 54.7 ± 16.6 indicating lower confidence (Figure 2G). Cloud genes thus appear to be enriched with low-quality gene models (short sequences, no InterPro domains, weaker structure models), which is further supported by peptide-based evidence, derived from experimental proteomics studies (further details below), where about 9.9% singleton pan-genes are supported by more than two peptides per gene. In contrast, 52.1% core RAP-DB genes show more than two peptides per gene as evidence for the pan-gene cluster (Figure 2H).

### Support for rice pan-proteome

To determine experimental support for the rice pan-genes, we performed large-scale re-analysis of public proteomics (mass spectrometry) data, against a comprehensive protein database derived from all possible gene models within the pan-gene set. We searched 19 public datasets (∼30 million MS/MS spectra), sourced and pooled from multiple rice varieties to maximize total coverage. All datasets were from “shotgun” methods, i.e. proteins were digested into a total peptide pool prior to LC-MS/MS. As a result, for peptides that can match to more than one gene product (per genome), it is not straightforward to determine which is the correct assignment. Using an all peptide-to-protein mapping, we demonstrate potential peptide-based evidence using >200,000 distinct peptides for 435,196 gene models and 293,844 genes present in 28,658 pan-genes.

Taking a parsimonious approach where a peptide can only be mapped to one protein, we found around 13K genes in each genome with at least two peptides, indicating strong evidence for the gene’s protein coding potential (Figure 3A). In agreement with the previous set of observations, we find a significantly higher number of peptides mapped to core and soft-core pan-genes than cloud pan-genes (Figure 3B). Out of all the peptides mapped to all the proteins, ∼73% (n = 168,151) of the peptides mapped to proteins in all 16 rice accessions indicating good support for these pan-genes, and that in general, based on current evidence, there is little deviation in sequences for readily detectable rice proteins across the pan-genome (Figure 3C).

**Figure 3.**
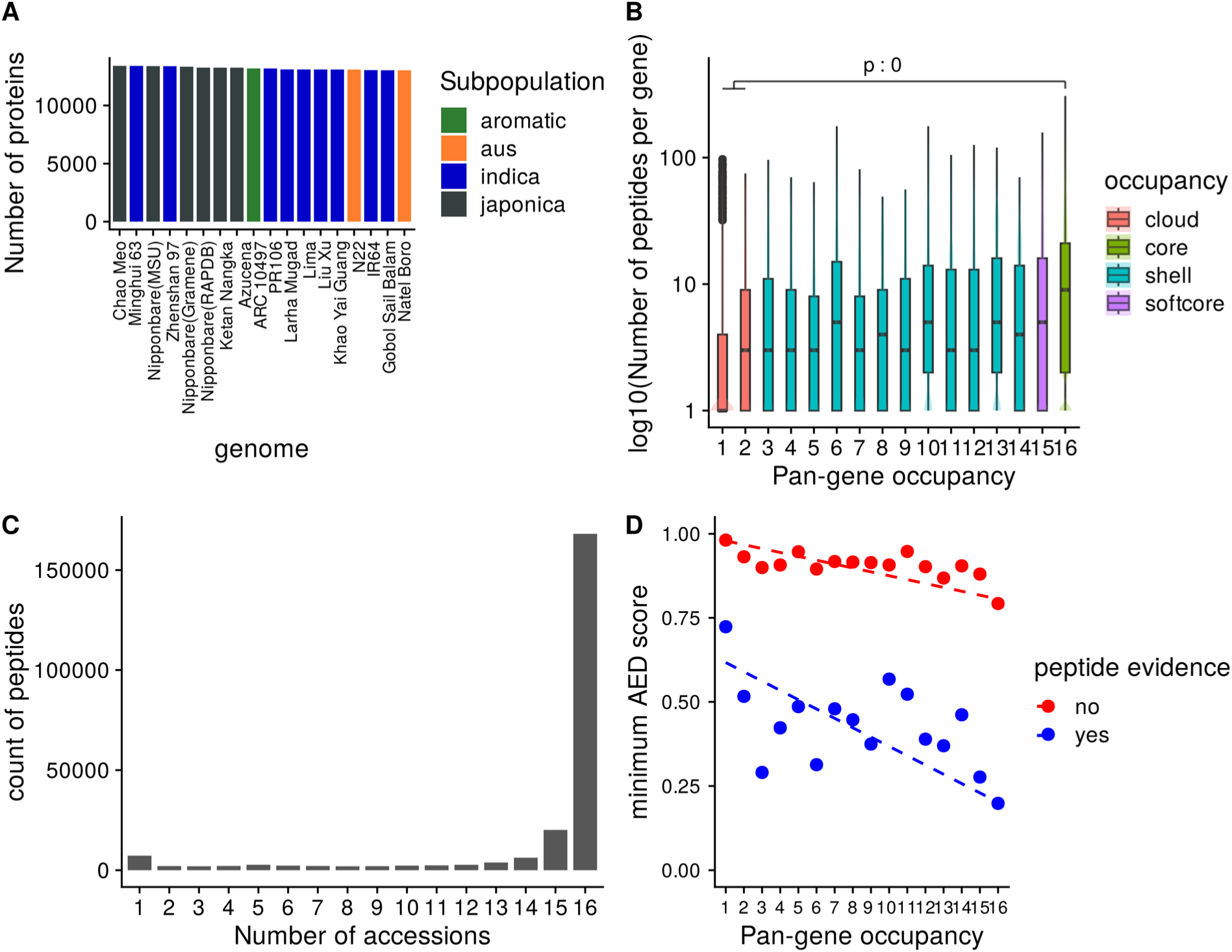
Experimental proteomics data support for pan-genes. A) Count of proteins per annotation set with at least two supporting peptides, B) log counts of peptide sequences versus genome occupancy, C) absolute counts of peptides mapped to proteins from 1-16 genomes, D) scatter plot of mean (min) AED score across three transcriptomics datasets for those (RAP-DB) gene products with or without peptide evidence, by genome occupancy of the containing pan-genes; dotted line = linear regression; indicating that gene products with peptide evidence have better (lower AED) support from transcriptomics data.

We further demonstrate the relationship between proteomics evidence (those genes with or without proteomics evidence) and AED score (derived from transcriptomes mapped to the RAP-DB gene representative of each pan-gene cluster) in Figure 3D. Proteins without experimental support tend to have high AED scores (ranging from 1 to 0.8, depending on genomes occupancy), whereas those with experimental support and high occupancy have low AED scores, indicative of well-supported, high-confidence gene models.

### Exploring the predicted protein domains of pan-genes

We ran InterProScan^15^ over all input proteins available within the pan-gene set, assigning Pfam^16^ and InterPro^17^ domains. For each pan-gene, we assigned additional datatypes and statistics, including the total number of different domains identified, the genomes (and proteins) containing those domains, and a representative domain for the cluster (most common across genomes), available in Supplementary table S5. We then assigned which proteins within a pan-gene cluster contain the representative domain, and which proteins are outliers i.e. do not contain the most common domain. Of the 42,501 pan-genes with at least two members, 26,044 have at least one assigned Pfam domain (and 28,507 at least one InterPro domain). Of these, 21,779 are “domain consistent” clusters (83%), having zero genomes lacking the most common Pfam domain. Similarly, 88% of pan-gene clusters have a consistent common InterPro domain. In Supplementary table S6 we show the counts of proteins that do not contain the most common InterPro domain per cluster per input genome for the 3485 pan-genes that are not “domain consistent”. There are around 400-600 proteins per input annotation set that do not match the most common domain. RAP-DB and MSU gene models have higher counts but have not been annotated using the same pipeline as other genomes, and thus are less consistent than OsNip models compared to other genomes, and we cannot straightforwardly conclude that the gene models from these sources are less reliable.

We identified the most enriched domains found in the set of core versus cloud pan-genes and observed some distinct patterns of domain types (Figure 4A). Core pan-genes are enriched in domains such as AP2/ERF, WD-40, Helix-loop-helix, Myb domain, C2H2 zinc finger and homeobox domain. These domains are commonly present in DNA-binding transcription factor proteins, suggesting their conserved nature among core pan-genes.

**Figure 4.**
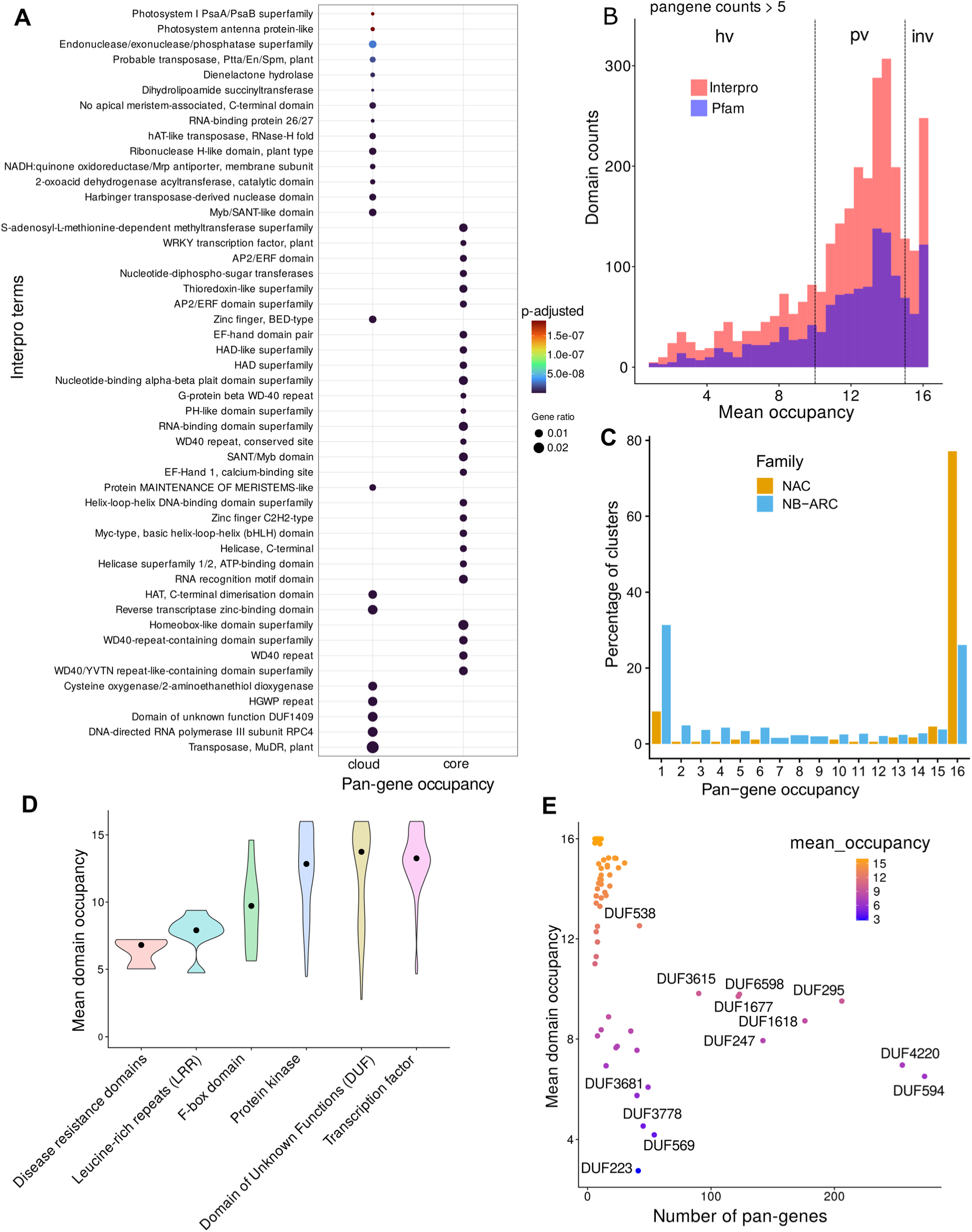
Exploration of predicted pan-gene domains. A) Enriched InterPro terms among cloud vs core pan-genes. Top 30 terms are shown in the plot. Size of the dot represents the enrichment factor and the color of the dot shows the p-value. B) Histogram shows the counts of domains within the pan-genes vs their mean domain occupancy for both Pfam and InterPro domains. The domains were filtered for presence in minimum 5 pan-genes. C) The plot shows the percentage of pan-gene clusters according to their occupancy status for two gene families, NAC domain containing TFs and NB-ARC domain containing genes. D) The variation in mean domain occupancy of various Pfam domains for the selected major gene groups is shown. The solid back dot shows the median value. E) The scatter plot shows the mean domain occupancy of various domains under the group ‘Domains with Unknown function’ (DUFs) vs the number of pan-genes with the respective domain. DUFs present in more than 50 pan-genes are highlighted in blue text.

Next, we explored the occupancy of protein domains across the pan-gene set. Variance in the distribution of domains can potentially inform on functional units that have either a tendency to become duplicated or have been more frequently introduced via introgression, compared to those that are invariable. Figure 4B contains a histogram of the mean occupancy of protein domains (InterPro and Pfam) showing a multimodal distribution. As noted in the methods, we then split the distribution into three groups: “invariable (occupancy) domains”, “partially variable” domains and “highly variable domains”. We hypothesize that gaining (or losing) copies of invariable domains are highly detrimental to rice. GO enrichment analysis of pan-genes containing these groupings of domains provide support to this hypothesis (Extended data figure 3). For example, pan-genes containing Ubiquitin-protein transferase and transcription factor DNA binding activity show conserved and invariable domains, whereas, domains involved in cell surface receptor signaling appear highly variable indicating the fast-evolving nature of these domains driven by selective breeding pressure, like disease resistance.

As a case study to test the pan-gene clusters, we analyzed two gene families (Figure 4C), i.e. the NAC transcription factor family (an invariable domain) and the NB-ARC disease resistance family (a highly variable domain)^18^. NAC transcription factor family genes have been characterized in the past for their roles in abiotic stress responses in rice^19^. We identified 175 NAC pan-genes with 142-154 genes in each genome, and the domain has a mean occupancy = 13.2. 77% of NAC pan-genes (n = 135) have core occupancy and 9% cloud (n=16) indicating a highly invariable domain (Figure 4C, Extended data figure 4). In comparison, the NB-ARC is a highly variable domain (mean occupancy = 7.2), identifiable within 1005 pan-genes with 432-518 genes per genome. NB-ARC family genes are mostly found with occupancy = 16 or 1, indicative of rapid introduction of additional copies or recent gene duplications in some genomes (Extended data figure 5). There is also a difference between NAC and NB-ARC in shell pan-genes: 9% of NAC pan-genes (n = 16) versus 34% of NB-ARC genes (n = 341). One could hypothesize that since NB-ARCs have a role in pathogen effector recognition, it is desirable for the plant to have a wide diversity of genes, but can tolerate the loss of a small number of genes in some genomes (leading to shell pan-genes), whereas loss of a transcription factor is highly detrimental, and the existence of cloud genes is explained by rare duplication events.

**Figure 5.**
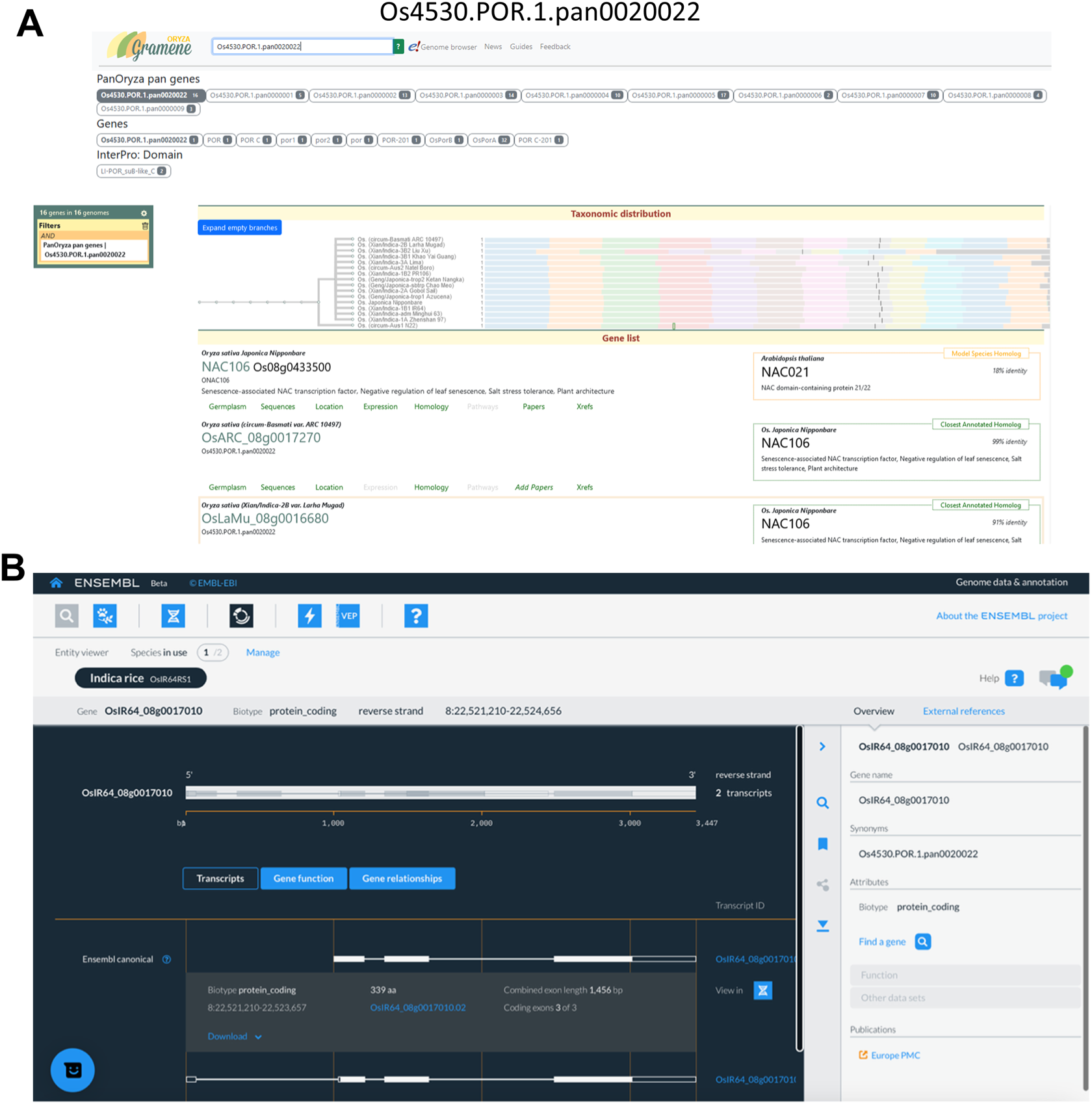
Representative example of infrastructure for pan-gene exploration using pan-gene Os4530.POR.1.pan0020022. A) The screen shot shows the record of pan-gene Os4530.POR.1.pan0020022 on the Gramene database (https://oryza.gramene.org/). B) The page shows gene record for a member gene OsARC_08g0017270 of Os4530.POR.1.pan0020022 on the Ensembl Beta Release.

A similar trend is seen with other domains like Leucine rich repeats (LRR, median occupancy of 7.50 ± 0.945), indicating the highly variable nature of these domains (Figure 4D). Given the role of LRRs in plant immune responses, the domain occupancy data are suggestive that these genes are fast evolving through breeding, for example where past gene duplications giving resistance to a given pathogen in cultivated rice have been selected. Notably, it is one of the domains associated with pan-genes enriched with highly variable GO term “cell surface receptor signaling pathway” (Extended data figure 3). Multiple domain types with the name ‘F-box’ show a partially variable occupancy (9.87 ± 2.66) compared to the least variable domains of transcription factor (13.3± 1.73) or protein kinase (13.4 ± 3.70) types. F-box domain containing proteins include those involved in the SCF type ubiquitin E3 ligase family, and in plants have multiple functional roles, including in stress responses. Our data also suggest past selective breeding pressure, giving rise to variability in the domain occupancy across the rice pan-genome.

Interestingly, the class of ‘Domains with Unknown function’ (DUFs) shows a median occupancy of 13.6 but with a wider interquartile range (± 5.49) with 46 domains present in more than 10 pan-genes (Figure 4E). Several DUFs (DUF594, DUF4220, DUF295, DUF1618, DUF6598 and DUF1668) are present in more than 100 pan-genes and have been classified as highly variable domains, which we believe may be indicative of past selection. The genes containing these domains are thus candidates for functional validation, and potential carriers of desirable traits.

### Infrastructure for pan-gene exploration

The pan-gene set has been loaded into popular, widely used databases, including Ensembl Plants and Gramene, and our own pan-gene resource (https://panoryza.org/). At Gramene (https://oryza.gramene.org/), it is possible to search with pan-gene identifiers, e.g. Os4530.POR.1.pan0020022, which is the pan-gene identifier for NAC106 (RAP-DB: Os08g0433500) (Figure 5A). NAC106 is a NAC transcription factor, which has been associated with salt tolerance, tiller angle and leaf senescence^20^. The Gramene search then returns the gene records within each of the genomes assigned to that pan-gene identifier. There are multiple views for exploring the data, including as phylogenetic trees within the set, or expanded to include orthologs from other plants. The MAGIC-16 assemblies and annotations have been added into Ensembl Plants and the new Ensembl site which is currently in beta (beta.ensembl.org). The pan-gene identifiers are visible as gene synonyms on beta, and will become available via Ensembl Plants in a future release (Figure 5B).

At the Pan-Oryza site, we have a JBrowse^21^ instance where each genome can be explored, with various tracks of data (Extended data Figure 6A). For the IRGSP reference, we have aligned gene models from other cultivars using LiftOff, enabling any variation in models from other genomes to be displayed. For NAC106, it is an occupancy=16 gene, with some suggested variation in exon 2 in N22 and Azucena varieties.

We have also created an R library, with code for interactive exploration of clusters using R Shiny through a web-browser. The application allows users to visualize the relative position of genes within a cluster across the 16 genomes, and explore the members and meta-data about each pan-gene (https://github.com/PGB-LIV/PanOryza-pan-genes-release-v1.0/tree/main/heatmap_app/) (Extended data Figure 6B).

A permanent DOI has also been created for the pan-gene matrix and identifiers (https://zenodo.org/records/14772953) so that they can be used now in other research projects. It is anticipated that they be updated in the future (e.g. yearly), under some of the following circumstances: i) gene models change; ii) additional genomes are added; and iii) there are refinements to the algorithm for creating pan-genes. Mappings will be made available, so that any users can track how and why pan-genes have changed. These resources can all be used by the rice research and breeding community now, to assist in mining the wide pool of genetic resources available for Asian rice.

## Discussion

New genomes are frequently being sequenced and assembled for many species, including rice. It is also reasonably straightforward to produce an annotation set (i.e. genes and protein sequences), although structural genome annotation remains very challenging to perform accurately^11^. The upshot is that archival repositories like INSDC contain an ever-increasing number of genome assemblies, and annotations of variable quality, and it is highly challenging for most research groups to interpret. For example, asking how a given gene varies in presence, absence, or expression across different rice varieties could not previously be performed without running complex, custom-built bioinformatics analyses.

Other groups have also explored or generated pan-genomes for rice. For example, Shang *et al.*^22^ sequenced, assembled and annotated a 251 accession panel, demonstrating as in our work, that LRR genes have variable copy number across genomes. Wu et al. generated a pan-genome from 74 weedy and cultivated rice accessions, building ortholog relationships using BLASTP, shedding light on genes involved in domestication. The Rice Gene Index (https://riceome.hzau.edu.cn) resource^23^ has also created ortholog groups across the RPRP, based on reciprocal best match method, creating >110,000 groups via this method. After filtering gene models from different input sets, our 78,000 pan-genes have reasonable agreement, with ∼22K pan-genes having identical membership to RGI clusters (Extended data figure 7).

Our work has a different focus from previous related efforts though, in that we have created a robust and stable pan-gene set for Asian rice, based on 16 very high-quality assemblies and gene annotation sets, supported by multiple types of experimental data. Persistent identifiers have been assigned to pan-genes, along with a system allowing for pan-genes to be updated over time, as source data or evidence changes, with the ultimate aim to make these resource straightforwardly available in end-user focused databases.

The availability of pan-genes in popular public databases allows for the rice research community to move now to working in the pan-genome context. For example, groups planning new genome wide association, gene expression, or proteomics studies can select to use the reference assembly (and gene/protein set) most closely matching their variety of interest, but using the pan-gene set created here to compare easily where SNPs/genes/proteins are present or absent across rice families, including in the reference genome IRGSP.

### Interpretation of pan-genes by their occupancy across genomes

Our QC metrics demonstrated that clusters are robustly formed, for example containing gene/protein sequences that are highly related across genomes, but with some outliers where sequences have low similarity. The clustering algorithm ensures that overlapping genes on different strands are not placed in the same cluster, but there are multiple other reasons why clusters (formed primarily based on gene span overlaps) contain only moderately similar or unrelated protein sequences. These include cases where one coding sequence has been predicted within introns, UTRs, or on a different frame from another gene model. In a few cases, the differences could be the result of true biological differences, but perhaps a default assumption should be that for incongruous pan-genes, some gene models are incorrect in one or more cultivars. The metrics we have generated allow such clusters to be identified, and work can begin to identify the correct coding sequence.

Exploring occupancy data points to several key conclusions. First, core pan-genes are well supported by experimental evidence (transcriptomics, proteomics) and have easily recognizable protein domains and structures. As a general trend, the extent and quality of experimental evidence rapidly decreases with occupancy – in some cases there are likely pseudogenes or only partial gene models that have been incorrectly predicted. In other cases, as demonstrated in the exploration of mean domain occupancy, genes under selective pressure have more variable occupancy.

### Implications for rice research and breeding

The concept of a reference genome remains a useful one, despite the wealth of genomes now available for rice (as with other species). It is a very costly and challenging exercise to annotate a genome sufficiently well i.e. performing manual or semi-automatic fixes to gene models, and performing functional annotation and validation of genes. However, it is evident that the IRGSP RefSeq does not fully represent the coding potential of the diverse set of varieties being grown around the world. In this pan-gene set, we find >5000 genes that are observed in at least two other *Oryza* subpopulations but absent in IRGSP. Another key finding is that by exploring the pan-gene set, we can see clear signatures of selection within protein domains. For example, some highly variable domains are associated with immune-related functions, and presumably resistant to pathogens and pests. Through identification of genes carrying currently under-studied domains (DUFs), and genes with recognizable domains that are present in only some subpopulations, these resources are ready for mining to find new alleles with desirable traits. Lastly, our approach for creating pan-genes and assigning globally unique (but easily updatable) stable identifiers, can be adopted by other communities attempting to deal with the complexity of multiple genomes and annotation sets within a species of interest.

## Methods

### Consistent gene annotation for the RPRP

To create the assembly of pan-gene sets across *Oryza sativa*, we first collected whole genome sequences, gene and protein sequences (all fasta format), and gene model coordinates (GFF format) for the RPRP rice population. We used as input to pan-gene set building gene models that have been generated by a single consistent pipeline, as described in^24^ for all 16 genomes. In brief, models were generated with MAKER-P, with *ab initio* gene prediction (SNAP^25^, Augustus^26^ and FGENESH^27^), deep RNA-Seq data (long and short read), and assignment of canonical gene models using TRaCE^12^. For the IRGSP (Nipponbare) reference, we created a merged annotation set from the three sources of gene models (RAP-DB; new models following the pipeline above, called OsNip; and MSU), using the following algorithm.

Protein coding genes from the three sets were merged based on gene overlap calculated using chromosomal coordinates and strand information. Genes were merged (protein-coding genes with CDS transcripts only) if the gene ranges overlapped by >=50% of the length of the shorter gene. The merged gene’s coordinates were adjusted to match the most extreme 5’ and 3’ coordinates of any protein-coding transcript within the merged gene. The name given to the merged gene was assigned from one of its members, giving priority to the RAP-DB identifier, followed by the OsNip identifier, and finally the MSU identifier. The MSU annotation set uniquely contain some genes tagged as transposable elements (TEs), which were not filtered at source (in case they had been incorrectly annotated as TEs), but for some analyses were later removed, as detailed below.

### Building pan-gene clusters and quality control

We ran the GET_PANGENES pipeline^10^ (version 11012024), which performs whole genome alignment using minimap2^28^ (version 2.17), followed by the use of bedtools intersect^29^ to determine overlaps in each pairwise alignment. All haploid gene models progressed to cluster building, except for the Liu Xu annotation set, where alternative heterozygous alleles of genes, which had been placed onto contigs as part of a diploid genome resolution process, were excluded from the pan-gene clustering step. Genome sequences and GFF3 files for the RPRP were sourced from Ensembl Plants (release 57, and unchanged in current release 60) and OsNip from Gramene.

GET_PANGENES was used to generate various quality control metrics, including running BUSCO (v 5.7.1), Clustal Omega^30^ (version 1.2.4) to generate protein-level multiple sequence alignments for each pan-gene cluster, followed by AliStat^31^ (version 1.14) to generate distance metrics. Core gene sets counts were generated using Tettelin^13^ and Willenbrock^14^ functions, fitted after 20 permutation experiments.

Pan-genes were classified as *core* if they contained a member from 100% of input genomes (all 16 in this case), *softcore* 95% of genomes (15 in this case), *cloud* (1 or 2 genomes) and *shell* (all other cases)^32^. Annotation Edit Distance (AED) scores were generated by TRaCE, using RNA-Seq data sourced from Zhou et al. ^24^. Each transcript was assigned an AED score, reflecting the proportion of the transcript overlapped by a StringTie model from each of the three RNA-Seq libraries.

Paralog and ortholog counts were generated from Ensembl Compara (build 111, https://ftp.ebi.ac.uk/ensemblgenomes/pub/plants/release-58/tsv/ensembl-compara/homologies). To generate this dataset, representative protein sequences of coding genes, from 116 plant genomes, were classified into clusters against a profile HMM library incorporating resources such as PANTHER^33^ and TreeFam^34^; with any remaining unclassified genes being included in a process of clustering by hcluster_sg (https://sourceforge.net/p/treesoft/code/HEAD/tree/trunk/hcluster/) using the results of a BLAST search^35^. Each cluster’s protein sequences were aligned by MCoffee^36^ or MAFFT^37^, and for the resulting alignments, TreeBeST (TreeFam) was used to generate a gene tree in which nodes were annotated as representing events such as a gene duplication or a speciation. Homology relationships were inferred from the annotated gene tree, with the type of homology between a given gene pair being determined by features of their evolutionary history, such as whether their last common ancestor represented a speciation or duplication event^38,39^. Source data is also available at https://plants.ensembl.org/info/genome/compara/index.html.

### Assignment of pseudo-position

For visualisation and sorting purposes, we assigned a *pseudo-position* for each pan-gene. We first assigned the mode (most common) chromosome from across the members within a given pan-gene. If the pan-gene has a member from Nipponbare matching the mode chromosome, we used the chromosome number and position from the Nipponbare gene as the *pseudo-position*. If the pan-genei does not has a member from the Nipponbare genomei, we scan through the members finding another genomex that contains a genexi, with a “proximal” genexj (following ordering of the pan-gene matrix by the mean chromosome position of all genes, within each mode chromosome), that is present in both genomex and Nipponbare genomei. We then calculate the pseudo_position = chr_pos_gene1j + chr_pos_genexi-chr_pos_genexj, using values of j from the list of relative ranked positions on the chromosome of genomex: (1,-1,2,-2,3,-3,4,-4,5,-5,6,-6,7,-7,8,-8). The final pseudo-position is taken for the lowest absolute value of chr_pos_genexi-chr_pos_genexj across all values tested.

The effect of this mapping is to estimate, approximately, if such a gene did exist on the Nipponbare chromosome, where would it be located. There are several reasons why this location is not necessarily an accurate reflection of the “true” position of the pan-gene, with regards to the Nipponbare reference, however it has a useful property that it maintains a sensible ordering of pan-genes for visualisation.

AlphaFold models for *Oryza sativa* (Reference id: UP000059680) were retrieved from AlphaFold protein structure database (https://alphafold.ebi.ac.uk/) and the mean prediction score calculated for each model. Gene expression data for 11,726 rice RNA-Seq samples was sourced from https://plantrnadb.com. Average FPKM values were calculated for each RAP-DB/MSU gene and assigned to the pre-computed pan-gene cluster.

### Assigning stable identifiers

We have designed an approach for assigning stable identifiers to pan-genes (following similar proposals made by https://www.agbiodata.org/), which may be repurposed for other plant genomes, as follows: [clade].[group].[version].panddddddd

clade = a two letter code for a species level pan-gene, or one letter code for genus level, followed by the NCBI Taxon ID i.e. Os4530 for *Oryza sativa*.

group = a unique three letter code for the consortium/group releasing the pan-gene set e.g. POR for our “PanOryza” consortium.

version = integers starting from 1, incremented for each new release.

panddddddd = pan-gene identifier, with digits 0000001 for pan-gene clusters with 2 or more members, and 1000001 for singleton clusters (containing only one gene).

The identifiers thus have the property that they can be globally unique, but can be produced by different consortia, using different sets of input genomes. We assigned pan-gene identifiers to all genes clustered as part of this pan-gene set. New releases could be created in the following circumstances: changes to genome assemblies or gene models, addition of new genomes or gene models, changes to the clustering algorithm.

### Generating a rice pan-proteome map

We next determined the experimental support for proteins in the rice pan-proteome, by performing large-scale re-analysis of public proteomics data. Datasets were sourced from the ProteomeXchange central repository^40^, using the inclusion criteria that a global (non-enriched) analysis had been performed on any rice tissue, sourced from any *Oryza sativa* variety, using a Thermo Fisher Scientific instrument (for simplicity of pipeline compatibility). Raw MS data were converted to mzML format^41^, using ThermoRawFileParser^42^. Datasets were searched with the parameters listed in Supplementary table S7, using the MSFragger search engine^43^ (version 3.5), against a protein database formed by merging the 16 protein sets from the RPRP gene models, as well as the RAP-DB (canonical and predicted proteins, version 2022-03-11) and MSU (version 7) protein sets. Common contaminants were added from the cRAP database (https://www.thegpm.org/crap/), and reversed decoy sequences added using the FragPipe utility. Peptide-spectrum matches from MSFragger in PepXML format^44^ were post-processed with the Trans-Proteomic Pipeline version 6^45^ (PeptideProphet^46^, iProphet^47^, ProteinProphet^48^). The 19 datasets that passed quality control (>500 proteins identified) were loaded into PeptideAtlas after thresholding the identifications such that the resulting protein-level FDR was 1%. The results of this process are available at https://peptideatlas.org/builds/rice/.

### Assignment of protein domains to pan-genes

Domains in the protein annotations of all 18 annotation sets (for 16 genomes) were identified using InterproScan-v5.62-94.0, and then mapped onto pan-genes and source gene models. We next assigned a representative domain for the cluster, as the most common protein domain found across genomes (in the case of more than one domain being tied “most common”, one is randomly selected). Using this most common protein domain (for both InterPro and Pfam) consistency of pan-gene clusters was checked to identify the genomes in which the representative domain is present.

Next, for every InterPro and Pfam domain, the “domain occupancy” was first calculated for each pan-gene across 16 genomes i.e. counting the number of genomes with a protein carrying the domain within the pan-gene (for IRGSP only OsNip assignments were included for this analysis to avoid biasing the Nipponbare domain counts). Next, “mean domain occupancy” was calculated as the mean of the domain occupancy across all pan-genes in which the domain can be found. Domains were filtered to include only those found in >= 5 pan-genes and were categorized based on their mean occupancy across pan-genes, as “highly variable” where mean occupancy < 10; “partially variable” for mean occupancy greater than 10 and less than 15; and “invariable” > 15. Interpro2Go annotations were used for identification of GO terms for pan-genes containing these three categories of domains. Top terms enriched within these pan-genes were identified using clusterProfiler (v4.6.2)^49^ with p-value cutoff <0.05, adjusted by the Benjamini-Hochberg method.

To identify the most significant InterPro domains in core versus cloud pan-genes we first filtered out clusters containing only transposable elements (TEs). Since regular protein-coding genes could be among the clusters which were labeled as TEs in past Nipponbare annotations (sourced from MSU), for pan-genes with occupancy = 1, we removed the cluster if it is annotated as a TE in the InterProScan results or the original MSU annotations. Further for pan-genes with occupancy greater than 1, a cluster was filtered out only if all the genes in a cluster are identified as TEs in the InterProScan results. With these pan-genes filtered for TEs, we used the enricher function of the clusterProfiler as above to identify the significant InterPro term in each occupancy class (p-value = 0.05, adjusted by Benjamini-Hochberg method). The redundant term descriptions within each occupancy class were collapsed using the stringdist function (method Jaro-Winkler distance). The top 30 terms for core and cloud occupancy class based on the adjusted P-value were used for the dotplot, where size and color of the dot shows the GeneRatio (term enrichment factor) and adjusted P-value, respectively.

### Analysis of gene families

For the NAC transcription factor (TF) gene family, query protein sequences from Plant TFDB^50^ were used for blastP (ncbi-blast, v2.12.0, e-value < 1E-05) and HMMER (HMMER 3.3, e-value < 1E-3) search against protein sequences from all gene models within the pan-genes. Results were checked for Pfam (PF02365) and InterPro domain IPR003441, IPR044799 or IPR036093, and pan-genes containing at least one protein with the domain were retained and labeled positive for ‘NAC’.

The NB-ARC domain (Nucleotide-Binding domain shared by Apaf-1, certain R-proteins, and CED-4) is a conserved nucleotide-binding domain found in a variety of proteins across different organisms, including plants, animals, and fungi. It plays a crucial role in processes like apoptosis (programmed cell death) and innate immune responses. The identification of NB-ARC domain-containing proteins in RPRP lines was carried out using the Gramene Oryza API^51^. A targeted query was employed to retrieve genes annotated with the InterProScan domain IPR002182 (NB-ARC) from gene trees. The query output, provided as a CSV file, was then filtered by the system_name field to isolate entries corresponding to RPRP genomes. These filtered genes were subsequently mapped to their respective pan-gene clusters and were designated “NB-ARC” if any pan-gene member from a genome was identified within the NB-ARC domain-containing gene set.

### Infrastructure development

The PanOryza site incorporated JBrowse (version 1.16.11). For display purposes, gene models from other genomes were mapped to the Nipponbare reference using LiftOff^52^ (v1.6.3). The interactive heatmap visualisation of pan-genes was created using R package InteractiveComplexHeatmap (Gu, et al., 2021). As noted in the Results, pan-genes identifiers have been loaded into Ensembl Plants and Gramene for querying. The protein sets derived from the RPRP set are also available within the UniProt Knowledge base^53^ (https://www.uniprot.org/proteomes?query=oryza+sativa).

### Comparison of pan-genes with RGI

We ran a comparison between the pan-genes in this work with the “Ortholog Gene Indices” (OGI clusters) from the Rice Gene Index (https://riceome.hzau.edu.cn). We first matched clusters that shared the same RAP-DB (Nipponbare) identifier, and excluded Minghui 63, Zhen Shan 97 and Gramene annotations as these were created using different set of identifiers / gene model annotations. For the remaining members, we calculated the percentage agreement of identifiers within each pan-gene. For this, the percentage similarity was calculated as matching counts / total counts where total counts is the number of unique identifiers in a combined set of OGI and pan-gene; matching counts is the number of identifiers matched between OGI and pan-gene.

## Code availability

Code for GET_PANGENES available from: https://github.com/Ensembl/plant-scripts/blob/master/pangenes/, code for recreating analyses in this manuscript available from https://github.com/PGB-LIV/PanOryza-pan-genes-release-v1.0.

## Acknowledgements

We also acknowledge the Center for Quantitative Life Sciences at Oregon State University for hosting our web servers.

## Extended data figures

**Extended data figure 1.**
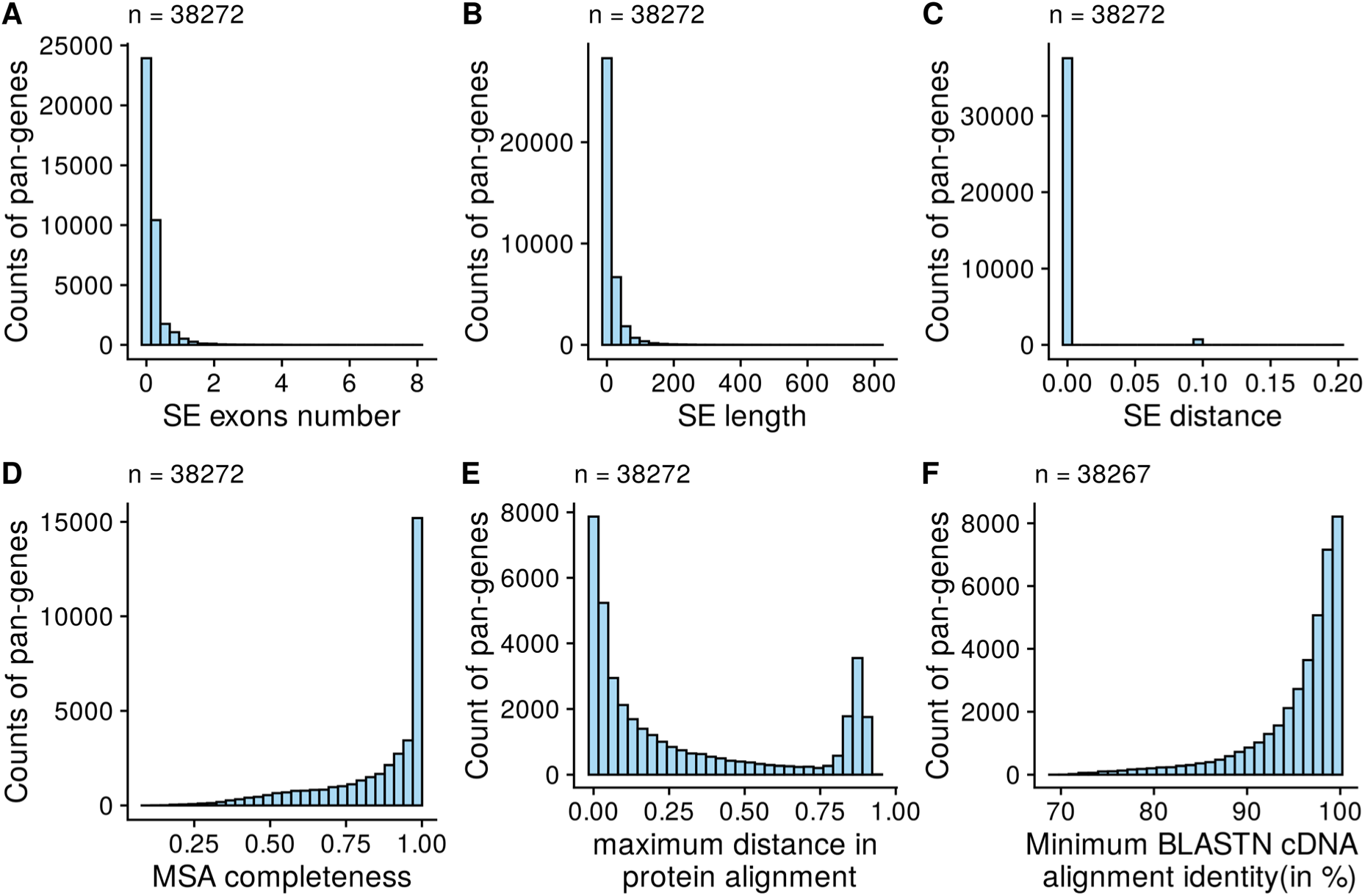
Quality control statistics of pan-genes. A) The standard error (SE) in the count of exons, B) SE of protein length, C) SE of sequence distance among all pan-gene members (transcripts/proteins) is plotted against the counts of pan-genes. D) The completeness of multiple sequence alignment (Ca values, calculated as in Wong et. al. 2020) and E) maximum sequence distance among pan-gene members is plotted against the counts of pan-genes. F) Minimum BLASTN cDNA alignment identity among members of pan-gene is shown against the number of pan-genes.

**Extended data figure 2.**
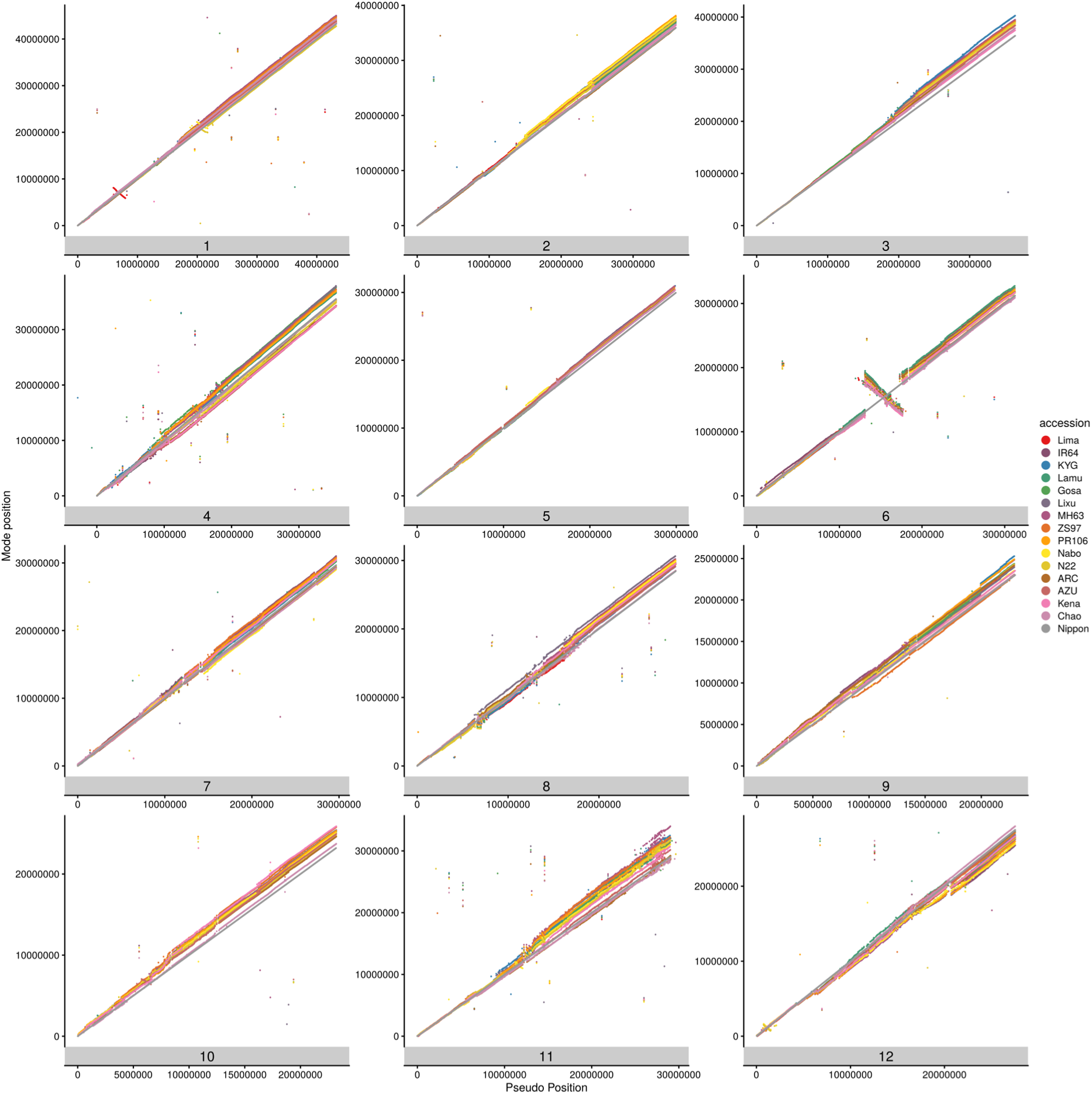
Position of genes on chromosomes across the RPRP accessions. Scatter plots show the positions of genes across the RPRP on each of the 12 rice chromosomes (y-axis) versus the position on the reference genome or inferred pseudo-position on the x-axis.

**Extended data Figure 3.**
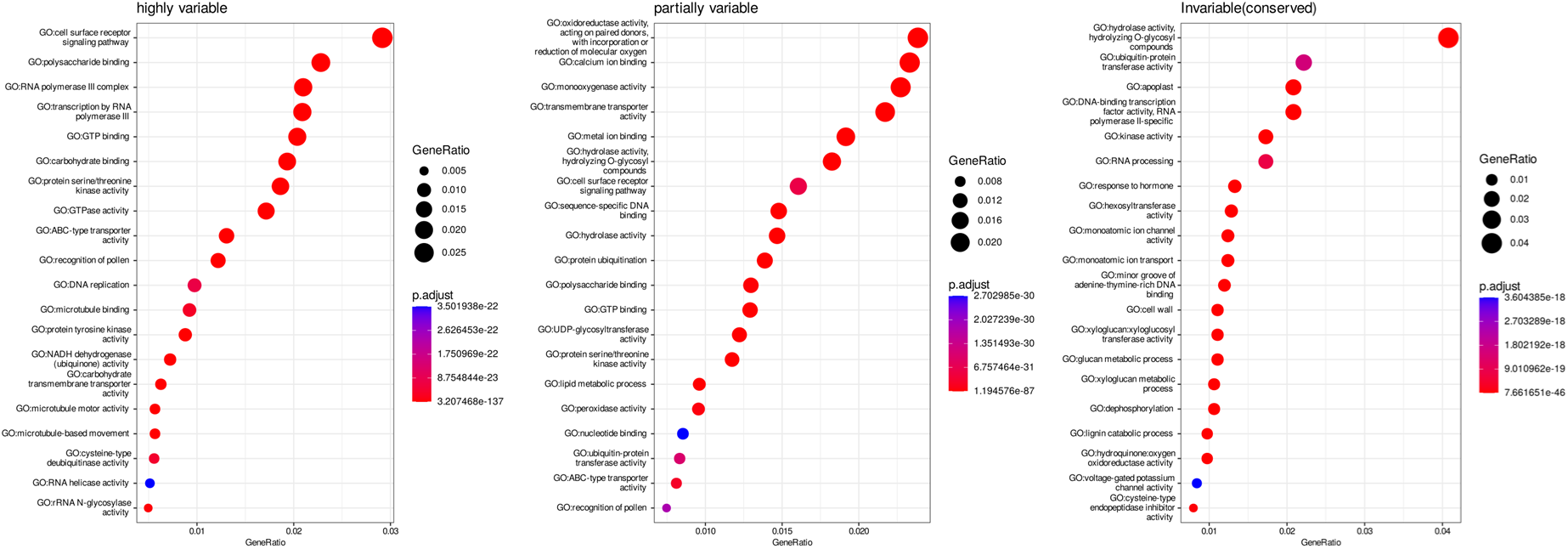
GO enrichment analysis for domains of variable occupancy. Most significant gene ontology terms for proteins containing domains classified as highly variable, partially variable or invariable.

**Extended data Figure 4.**
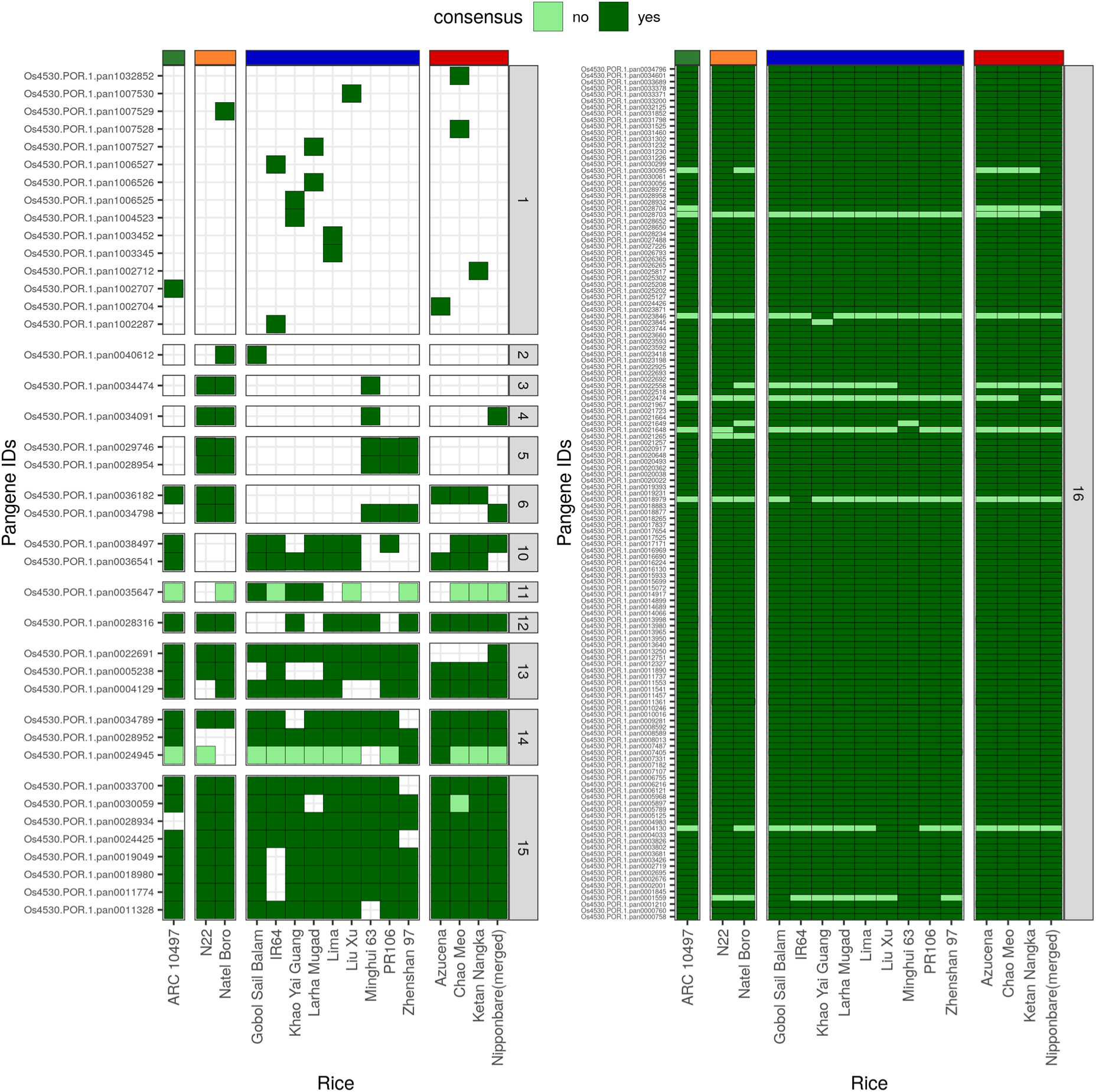
Pan-genes of NAC transcription factor family. The heatmap shows the pan-genes with at least one member protein containing the NAC domain. The vertical bars coloured grey on the right of heatmap show the occupancy of the pan-gene indicated by a number inside the box. The horizontal bars on top of the heatmap represents the varietal group colored red for *japonica*, blue for *indica*, orange for *aus* and green for *aromatic* rice accessions. The consensus presence /absence of the domain in a family member protein is indicated by dark and light shades of green in the box, respectively, whereas absence of the gene member is indicated by white box.

**Extended data Figure 5.**
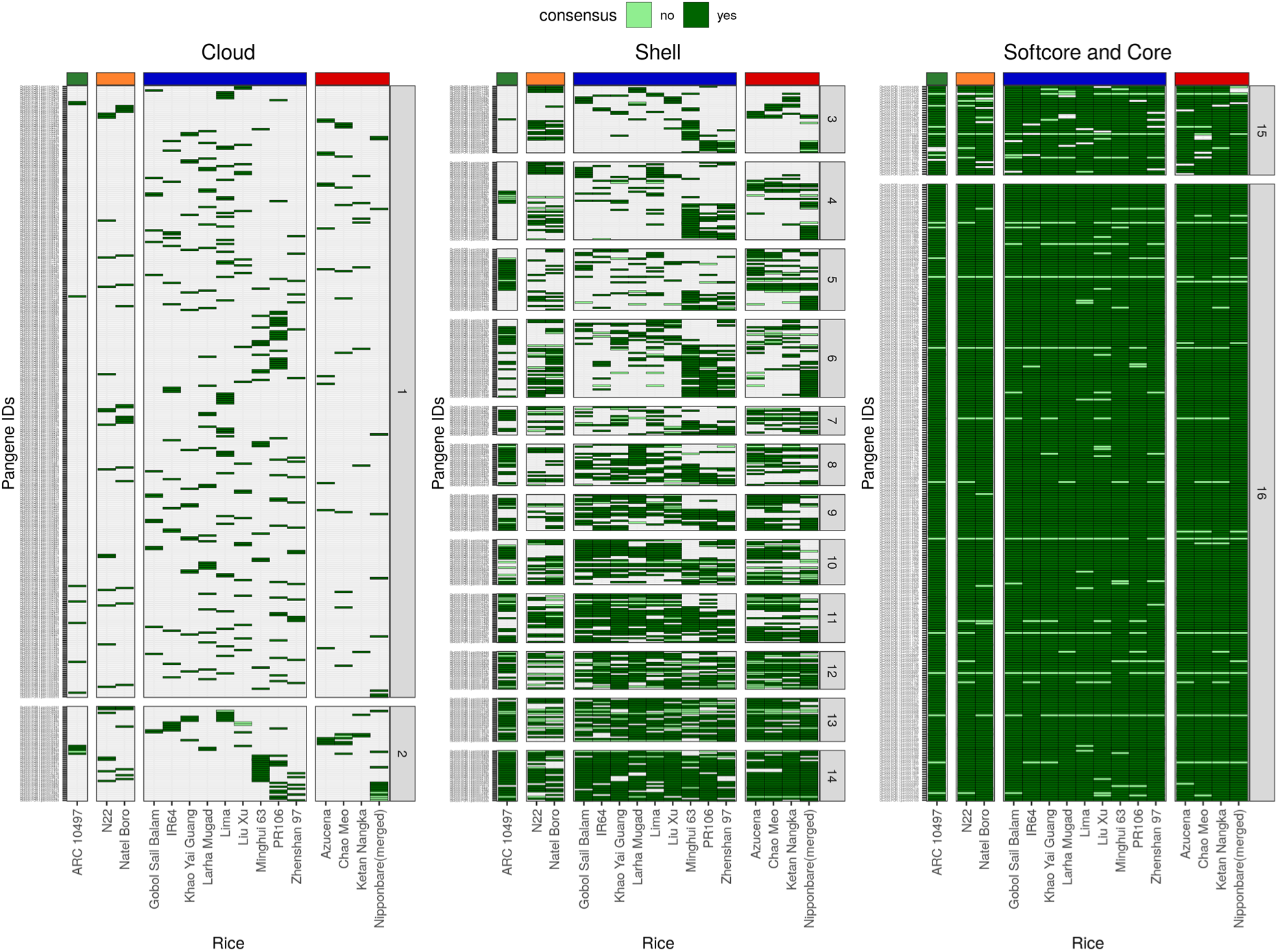
Pan-genes of NB-ARC family in rice. The heatmaps show pan-genes with at least one member protein containing the NB-ARC domain. The three heatmaps are arranged according to the occupancy classes – cloud, shell and softcore and core. The vertical bars coloured grey on the right of heatmap show the occupancy of the pan-gene indicated by a number inside the box. The horizontal bars on top of the heatmap represents the varietal group colored red for *japonica*, blue for *indica*, orange for *aus* and green for *aromatic* rice accessions. The consensus presence /absence of the domain in a family member protein is indicated by dark and light shades of green in the box, respectively, whereas absence of the gene member is indicated by white box.

**Extended data Figure 6.**
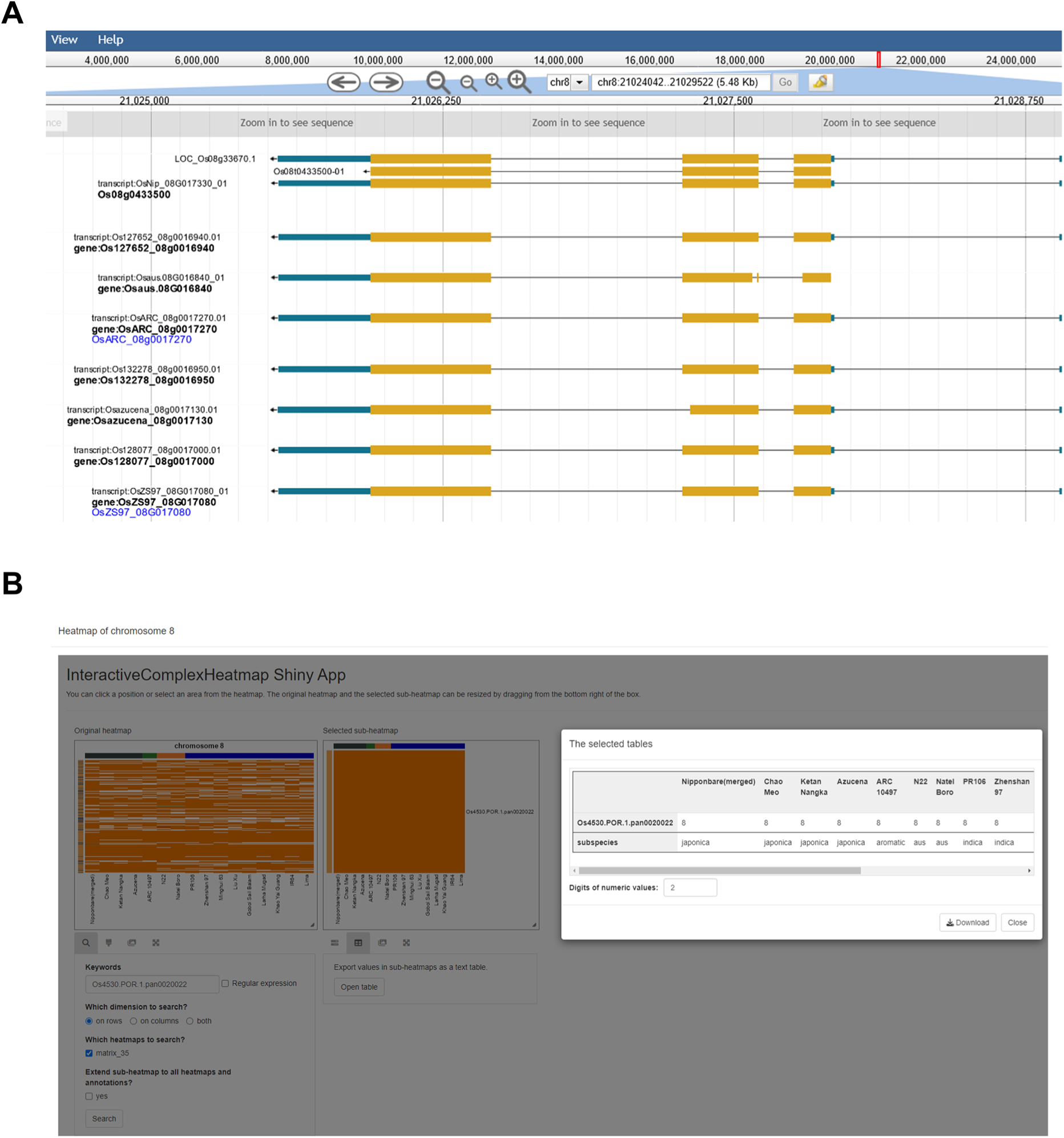
Representative examples of infrastructure for pan-gene exploration. A) The screen shot shows the record of pan-gene Os4530.POR.1.pan0020022 on the Gramene database (https://oryza.gramene.org/). B) A Rice Genome Browser (Jbrowse) visualisation of the Os4530.POR.1.pan0020022 on PanOryza project website (https://panoryza.org/). B) The panel shows the an interactive heatmap of chromosome 8 where all members of Os4530.POR.1.pan0020022 are located (detailed on the sub-heatmap and table). The pan-gene interactive heatmap for all chromosomes has been made available as a shiny application (https://github.com/PGB-LIV/PanOryza-pan-genes-release-v1.0/tree/main/heatmap_app).

**Extended data Figure 7.**
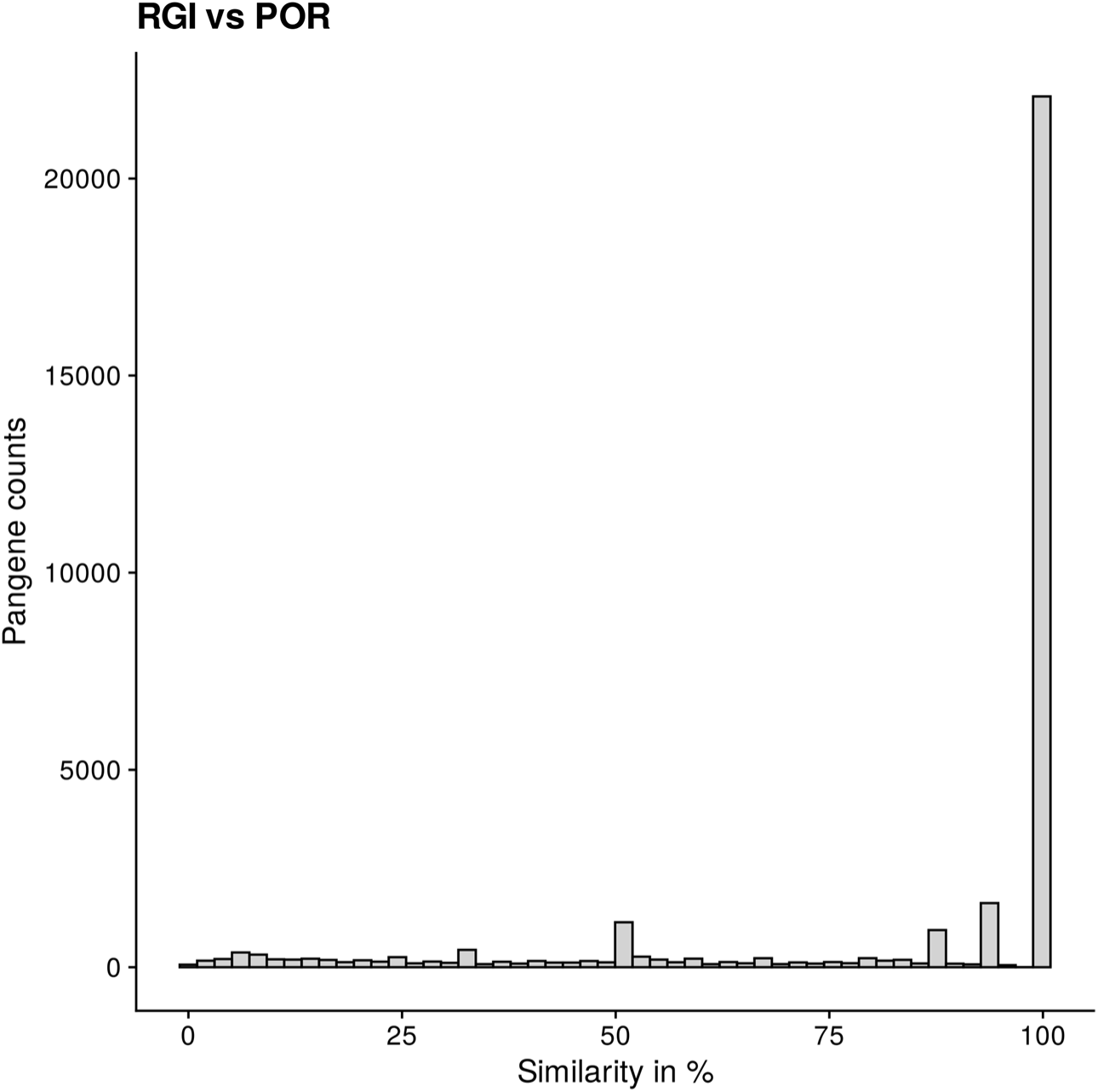
Comparison of current pan-genes with rice gene index. The plot shows percentage agreement between the pan-genes in this work with the OGI clusters from the Rice Gene Index (excluding Minghui 63, Zhen Shan 97 and Gramene annotations). The percentage agreement of identifiers within each pan-gene is indicated on x-axis and the counts of pan-genes on the y-axis.

## Source Data

The input data for running GET_PANGENES pipeline and the output files generated by the pipeline have been deposited at zenodo (https://zenodo.org/records/14772953).

## Supplementary Information

Supplementary table S1. Matrix of pan-genes containing transcript identifiers.

Supplementary table S2. Quality control statistics of pan-genes.

Supplementary table S3. Positions of genes within pan-gene clusters and pseudo-positions of pan-genes.

Supplementary table S4. Proportions of genomes of varietal groups - *Japonica*, *Aromatic*, *Aus*, *Indica* within each pan-gene.

Supplementary table S5. Summary of protein domains of pan-genes and respective representative domain information.

Supplementary table S6. Inconsistent pan-genes with member proteins lacking the most common InterPro or Pfam domain.

Supplementary table S7. Parameters used to generate the rice pan-proteome map.

